# RB1 loss overrides PARP inhibitor sensitivity driven by RNASEH2B loss in prostate cancer

**DOI:** 10.1101/2021.11.08.467839

**Authors:** Chenkui Miao, Takuya Tsujino, Tomoaki Takai, Fu Gui, Takeshi Tsutsumi, Zsofia Sztupinszki, Zoltan Szallasi, Kent W. Mouw, Lee Zou, Adam S. Kibel, Li Jia

## Abstract

Current targeted cancer therapies are largely guided by mutations of a single gene, which overlooks concurrent genomic alterations. Here, we show that RNASEH2B, RB1, and BRCA2, three closely located genes on chromosome 13q, are frequently deleted in prostate cancer individually or jointly. Loss of RNASEH2B confers cancer cells sensitivity to poly(ADP–ribose) polymerase (PARP) inhibition due to impaired ribonucleotide excision repair and PARP trapping. When co-deleted with RB1, however, cells lose their sensitivity, in part, through E2F1-induced BRCA2 expression, thereby enhancing homologous recombination repair capacity. Nevertheless, loss of BRCA2 re-sensitizes RNASEH2B/RB1 co-deleted cells to PARP inhibition. Our results may explain some of the disparate clinical results from PARP inhibition due to interaction between multiple genomic alterations and support a comprehensive genomic testing to determine who may benefit from PARP inhibition. Finally, we show that ATR inhibition can disrupt E2F1-induced BRCA2 expression and overcome PARP inhibitor resistance caused by RB1 loss.

## Introduction

Alterations of DNA damage response and repair (DDR) are associated with genomic instability, a hallmark of cancer, including prostate cancer (PCa). Genomic studies have revealed that about 10% primary and 27% metastatic PCa tumors have genomic loss (mutation or deletion) of at least one gene involved in DDR with BRCA2 being the most frequently mutated gene (*1, 2*). These alterations have been correlated with therapeutic vulnerabilities in PCa cells. Specifically, defects in homologous recombination repair (HRR) would predict the response to Poly (ADP-ribose) polymerase (PARP) inhibition. PARP is a family of enzyme involved in various cellular processes, notably DNA repair and genomic stability. PARP inhibitors (PARPis) are a new type of targeted therapy, which works by preventing PARP1 and PARP2 from repairing DNA single-strand breaks and resulting in stalled replication fork by trapping PARP1 and PARP2 on the DNA breaks (*3, 4*). These effects contribute to accumulation of DNA double-strand breaks (DSBs) that HRR-deficient cells cannot repair efficiently, causing overwhelming DNA damage and apoptotic cell death. BRCA1 and BRCA2 (BRCA1/2) encode proteins essential for HRR. Cancer cells lacking BRCA1/2 depend instead on PARP-regulated DNA repair and are hypersensitive to PARP inhibition (*5, 6*). Four PARPis (olaparib, NCT02987543; rucaparib, NCT02975934; niraparib, NCT02854436; and talazoparib, NCT03148795) are under clinical investigation in PCa, leading to regulatory approvals of olaparib and rucaparib for the treatment of metastatic castration-resistant prostate cancer (CRPC) patients with HRR deficiencies or deleterious BRCA1/2 mutations (*7–12*). While the results from these clinical trials have shown that patients with tumors harboring BRCA1/2 mutations benefit from PARP inhibition with high response rate, the degree to which patients with non-BCRA genomic alterations respond to PARPis remains unclear after gene-by-gene analysis.

To expand the efficacy of PARPis to tumors with non-BRCA alterations, efforts have been made to find new vulnerabilities for PARP inhibition in different cell models. Clustered regularly interspersed short palindromic repeat (CRISPR)/Cas9 loss-of-function genetic screen is a powerful approach to identify genes, once deleted, make cells more sensitive to PARP inhibition. Using this approach, recent studies have discovered that inactivation of enzymes involved in excision of genomic ribonucleotides or aberrant nucleotides may create vulnerability of cancer cells to PARP trapping (*13, 14*). The alteration of genes encoding these enzymes are potential genomic biomarkers or actionable targets for PARPis. RNASEH2B is one of these genes, which is particularly intriguing for PCa because it’s frequently deleted in primary and metastatic PCa tumors. RNASEH2B protein is one of the three subunits comprising ribonuclease (RNase) H2 complex that cleaves the RNA strand of RNA:DNA heteroduplexes, as well as single ribonucleotides embedded in DNA and play a role in DNA replication (*15*). It has been reported that inactivation of RNase H2 confers sensitivity to olaparib due to its function in ribonucleotide excision repair, loss of which leads to PARP trapping on DNA lesions (*13*). However, after investigating the publicly available PCa genomic datasets, we have observed that RNASEH2B is commonly co-deleted with two physically close genes RB1 and BRCA2. While deletion of RNASEH2B may confer PCa cells sensitive to PARP inhibition, the response may vary when RB1 and BRCA2 are co-deleted.

Targeted cancer therapies are increasingly being guided by genomic sequencing. However, current genomically-driven clinical decision making is largely based on mutations of a single gene. The potential impact of concurrent genomic alterations on therapeutic response has been overlooked. We speculate that combinatorial effects of compound genomic alterations may sway the synthetic lethality of a single gene deletion with PARP inhibition. Here, we investigate PARPi response of PCa cells after RNASEH2B deletion and co-deletion with RB1 and BRCA2 in preclinical models. Our study demonstrate that concurrent genomic deletions may have opposing impacts on PARPi response, supporting the utility of a comprehensive genomic testing instead of a single gene-based prediction in future clinical practice.

## Results

### Compound deletions of RNASEH2B, RB1, and BRCA2 genes in PCa

To determine genes associated with PARPi response, we analyzed five publicly available datasets of genome-wide CRISPR/Cas9 screens under the treatment with PARPi olaparib in hTERT-RPE1, HELA and SUM cells (*13, 16, 17*). We found a total of 79 genes common in at least two screens (Fig. 1A; table S1), loss of which sensitize cells to olaparib. We analyzed these genes for Gene Ontology (GO) term enrichment using a web-based gene annotation tool, DAVID 6.8 (*18, 19*). As expected, DNA repair processes were over-represented with “double-strand break via homologous recombination” being the most significantly enriched function (Fig. 1B). Out of this list, 13 genes are common to four screens, among which RNASEH2B is the most frequently deleted in both primary (17% homozygous deletion) and metastatic (12% homozygous deletion) PCa tumors (Fig. 1C), followed by FANCA, ATM, and BRCA1 in primary tumors and ATM, RAD51B, and BARD1 in metastatic tumors, respectively. The frequency of these genomic alterations was markedly increased when heterozygous deletions were counted as well (fig. S1A). The proteins encoded by RNASEH2A, 2B and 2C are three subunits of the RNase H2 enzyme complex (*20*). Deletion of any single subunit sensitizes cells to olaparib due to impaired RNase H2 function in ribonucleotide excision repair creating PARP-trapping lesions (*13*). While all three subunits are required for the function of the RNase H2 enzyme, the prevalence of RNASEH2B deletion make it an attractive biomarker to predict PARPi response in PCa.

RNASEH2B resides on chromosome13q14, which is a genomic region with frequent focal and arm-level deletion or loss of heterozygosity in PCa (*21–23*). In primary PCa tumors from the TCGA cohort (*24, 25*), we found that RNASEH2B is constantly co-deleted with a well-known tumor suppressor gene RB1 proximally located within a distance of 2.5 Mb (Fig. 1D; fig. S1B). In a small fraction of tumors, RNASEH2B and RB1 are co-deleted together with BRCA2, which is located about 18.5 Mb from RNASEH2B on chromosome 13q. In metastatic PCa tumors from the SU2C/PCF cohort (*26*), compound genomic alterations comprise single deletions and double/triple co-deletions of these three genes. In addition, we observed a positive correlation of the copy number values between these three genes in the TCGA cohort (fig. S2). An almost perfect correlation between RNASEH2B and RB1 genes indicated a potential focal deletion on chromosome13q14. Furthermore, we found that tumors with RNASEH2B and RB1 heterozygous or homozygous deletions exhibit significantly lower transcript levels in comparison with the wild-type tumors (Fig. 1E). Interestingly, the decrease was not observed in tumors with BRCA2 deletion, indicating more complex transcriptional regulation at the BRCA2 locus.

**Fig.1.**
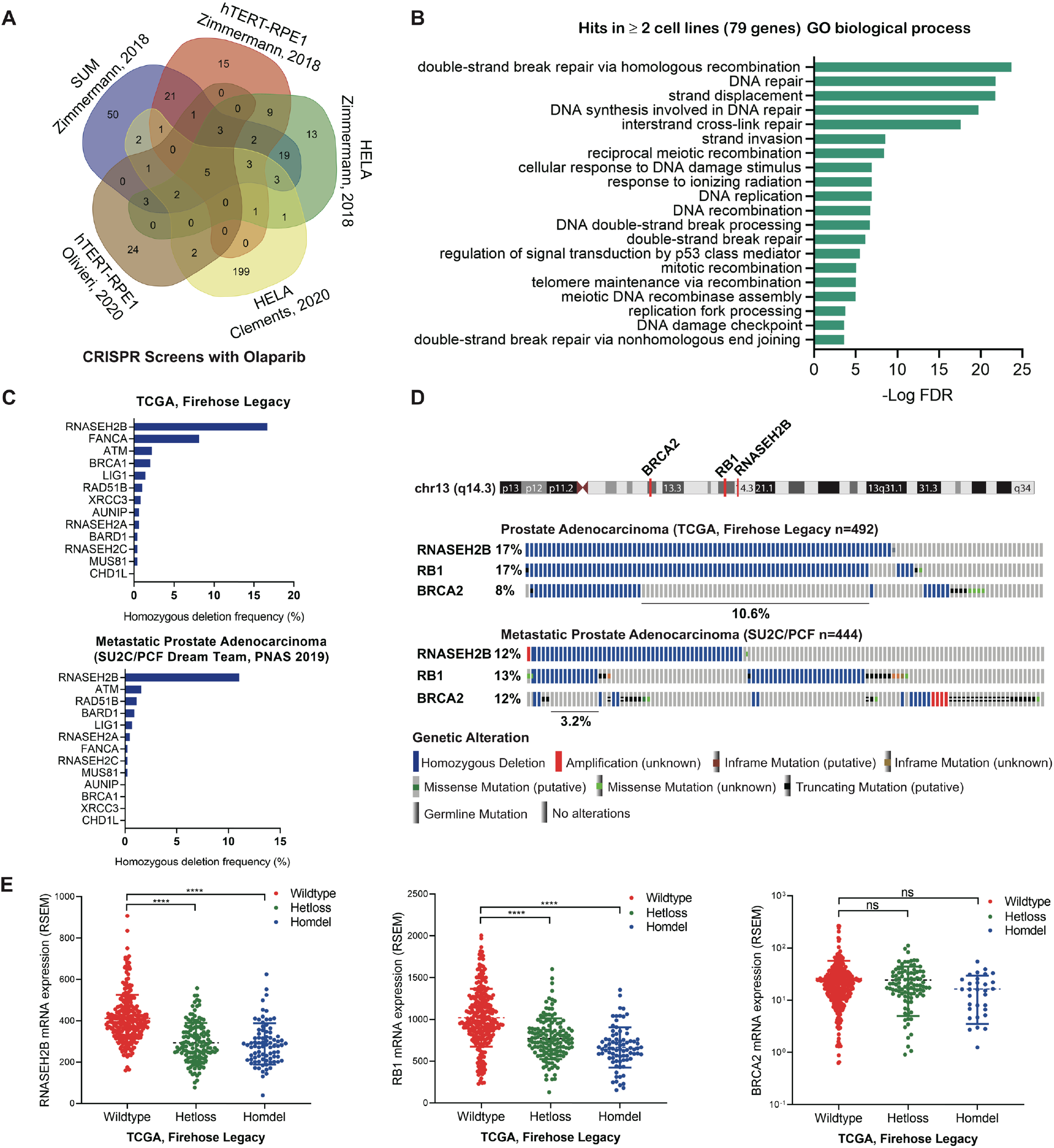
Identification of RNASEH2B loss as a potential biomarker to predict PARPi response in PCa. (A) Venn diagram showing the overlap between identified genes from five CRISPR/Cas9 screens with olaparib treatment. (B) Gene Ontology (GO) terms enriched among identified genes common in at least two CRISPR/Cas9 screens. (C) Homozygous deletion frequency of 13 identified genes common in at least four CRISPR/Cas9 screens in primary (in the TCGA cohort) and metastatic (in the SU2C/PCF cohort) PCa tumors. (D) Genomic alterations of RNASEH2B, RB1 and BRCA2 genes on chromosome 13q in primary (in the TCGA cohort) and metastatic (in the SU2C/PCF cohort) PCa tumors. The RNASEH2B/RB1 co-deletion accounts for 10.6% and 3.2% of cases in each cohort respectively. (E) The mRNA levels of RNASEH2B, RB1 and BRCA2 in primary PCa tumors (in the TCGA cohort) harboring wildtype RNASEH2B, heterozygous (Hetloss) and homozygous (Homdel) RNASEH2B deletions. *P*-values determined by two-tailed *t* test. **** *p*<0.0001 and not significant (ns).

### Deletion of RNASEH2B renders PCa cells sensitive to PARP inhibition

While previous studies have demonstrated that RNASEH2B genetic deletion sensitizes cells to PARP inhibition (*13*), to what extent loss of RNASEH2B increases PARPi response in PCa remain unclear. Using CRISPR/Cas9 gene editing, we deleted RNASEH2B in PCa cell lines LNCaP, C42B, 22RV1, DU145, and PC-3. Two different single guide RNAs (sgRNAs) were used for RNASEH2B knockout (KO) in each cell line, while sgRNAs against adeno-associated virus integration site 1 (AAVS1) were used to generate corresponding control cell lines. RNASEH2B deletion was confirmed by Western blot (Fig. 2A). Genetic deletion of RNASEH2B significantly increased cell sensitivity to olaparib across all five cell lines, more so in androgen receptor (AR)-positive LNCaP, C42B and 22RV1 cells in contrast to AR-negative PC-3 and DU145 cells. The sensitivity was assessed by the half-maximal inhibitory concentration (IC50) (table S2). We observed 253-, 30-, and 103-fold change in LNCaP, C4-2B, and 22RV1 cells in contrast to 3- and 8-fold change in PC-3 and DU145 cell, respectively. Increased sensitivity to olaparib after RNASEH2B deletion is comparable to that after BRCA2 deletion in C4-2B cells (fig. S3), indicating a similar impact of both genes on PARPi response. Previous studies have shown that loss of RNASEH2B creates more DNA lesions for PARP trapping (*13*). We next examined PARP1 protein levels in both nuclear soluble and chromatin fractions after olaparib treatment. We observed increased PARP1 protein trapped onto the chromatin in RNASEH2B-KO C4-2B and 22RV1 cells compared to AAVS1 control cells (Fig. 2B). Furthermore, we found that RNASEH2B-KO cells were also sensitive to PARPis rucaparib and talazoparib [with strong trapping ability (*3, 27*)], but to a lesser extent, to veliparib (with poor trapping ability) (fig. S4). These results suggest that PARP-trapping ability is critical for PARPi-mediated cell death in PCa cells with RNASEH2B deletion.

**Fig.2.**
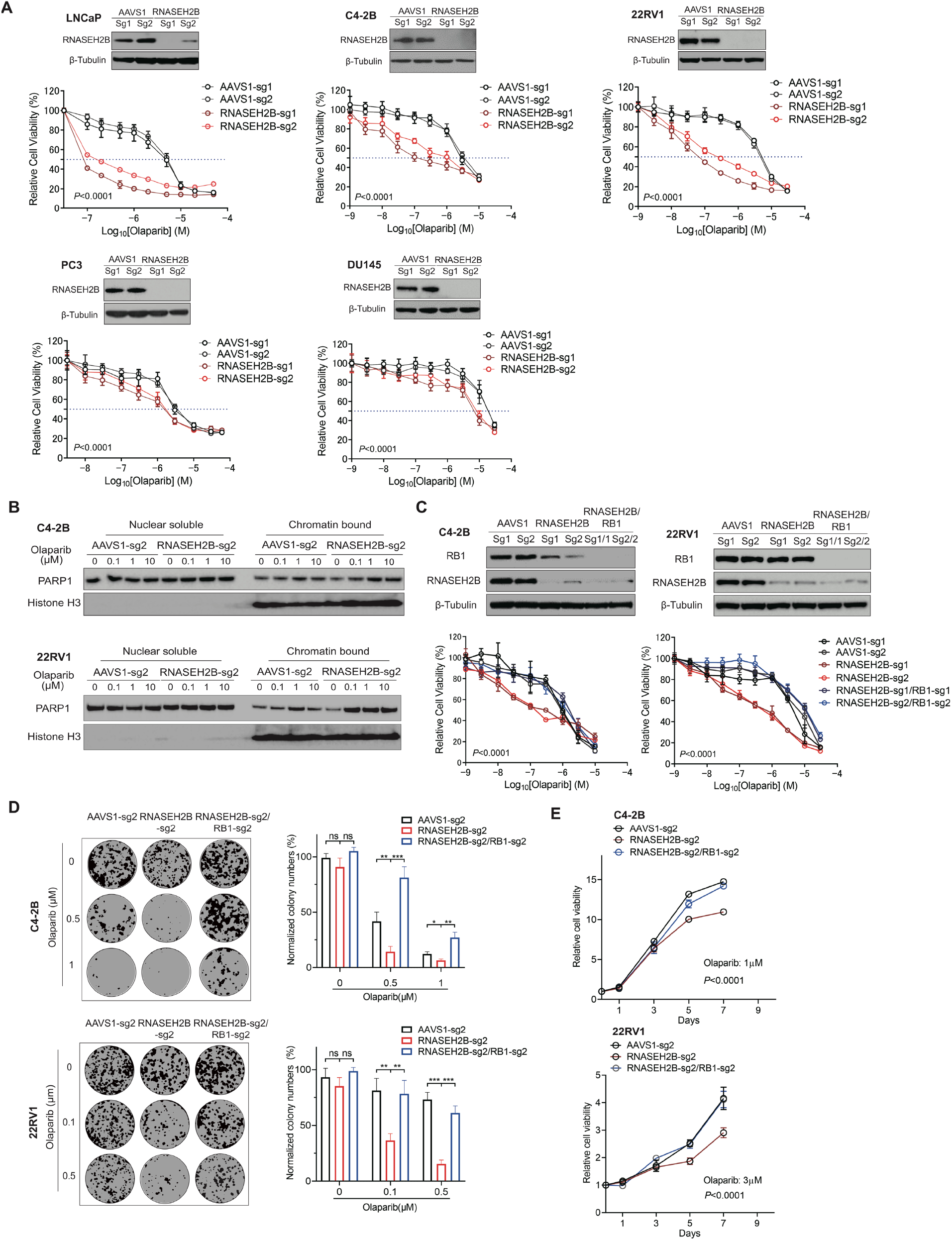
Impacts of RNASEH2B deletion or RNASEH2B/RB1 co-deletion on PCa cell response to PARP inhibition. (A) PCa LNCaP, C42B, 22RV1, DU145, and PC-3 cells, harboring two different sgRNAs against AAVS1 control (sg1 and sg2) and RNASEH2B (sg1 and sg2) were treated for 7 days with the indicated doses of olaparib. Cell viability was determined using alamarBlue assay (mean ± SD; n=3). Western blot showing RNASEH2B and BRCA2 protein levels in knockout (KO) and control cells. β-tubulin served as a loading control. (B) The protein level of PARP1 in nuclear soluble and chromatin-bound fractions of RNASEH2B-KO and AAVS1 control cells were determined by Western blot. (C) RNASEH2B single gene KO (SKO) RNASEH2B/RB1 double gene KO (DKO) C4-2B and 22RV1cells were treated for 7 days with the indicated doses of olaparib. Cell viability was determined using alamarBlue assay (mean ± SD; n=3). Western blot showing RNASEH2B and RB1 protein levels in AAVS1 control, SKO, and DKO cells. β-tubulin served as a loading control. (D) The growth of AAVS1 control, SKO, DKO C4-2B and 22RV1 cells were determined using colony formation assay after olaparib treatment for 14 days. (E) AAVS1 control, SKO, DKO C4-2B and 22RV1 cells were treated with olaparib for the indicated days. Cell proliferation rates were determined using alamarBlue assay. *P*-values determined by two-tailed *t* test or two-way ANOVA. * *p*<0.05, ** *p*<0.01, *** *p*<0.001 and not significant (ns).

### Loss of RB1 diminishes the sensitivity of RNASEH2B-deleted PCa cells to PARP inhibition

To determine whether co-deletion of RNASEH2B and RB1 impacts PARPi response, we deleted the RB1 gene in RNASEH2B single gene KO (SKO) C4-2B and 22RV1 cells to generate RNASEH2B/RB1 double gene KO (DKO) cells (Fig. 2C). We found that co-deletion of RB1 significantly reduced cell sensitivity to olaparib. In both cell viability and colony formation assays (Fig 2, C and D), PARPi sensitivity of RNASEH2B SKO cells was completed abolished by concurrent RB1 deletion. DKO cells showed significantly increased proliferation rate in comparison to SKO cells under olaparib treatment (Fig. 2E). It should be noted that deletion of RB1 alone reduced PARPi sensitivity in C4-2B cells (fig. S5), suggesting a potential intrinsic PARPi resistant mechanism rising from RB1 loss.

Using immunofluorescence staining for γ-H2AX foci, a marker for DNA DSBs, we detected significantly increased DNA damage in the nucleus of SKO C4-2B and 22RV1 cells after olaparib treatment for 24 hours compared to their corresponding control cells, which was not observed in DKO cells (Fig. 3A). Higher γ-H2AX protein levels were also detected in SKO cells after olaparib treatment using Western blot (Fig. 3B). Accordingly, the cleaved-PARP was increased in SKO cells compared to AAVS1 control and DKO cells, indicating undergoing apoptosis in RNAEH2B-deleted cells after olaparib treatment. Increased SKO cell apoptosis was further confirmed using Caspase3/7 activity assay (Fig. 3C).

**Fig.3.**
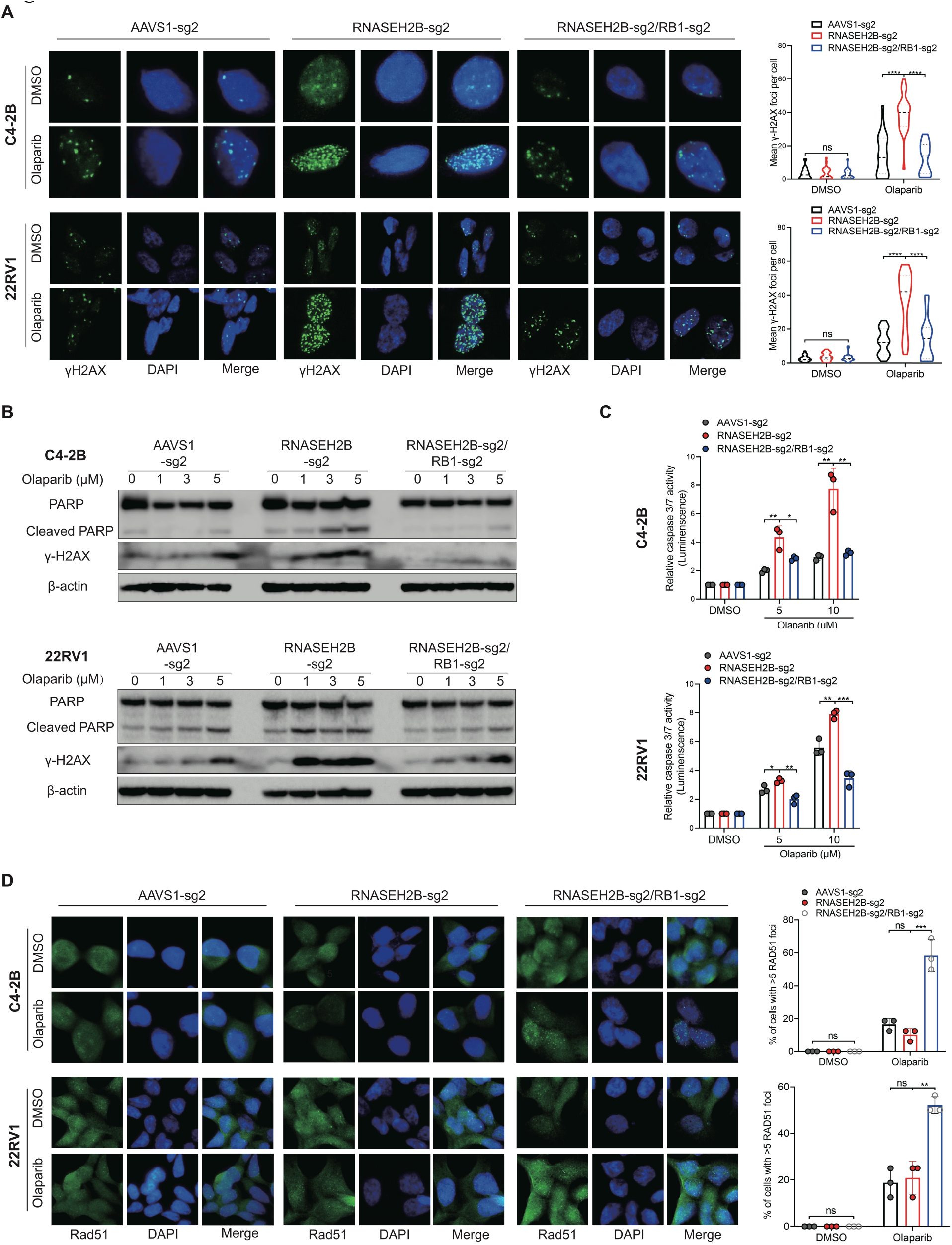
Impacts of RNASEH2B deletion or RNASEH2B/RB1 co-deletion on DNA damage, apoptotic cell death, and HRR function in PCa cells. (A) representative images of immunofluorescence staining for γ-H2AX foci in AAVS1 control, SKO, and DKO C4-2B and 22RV1 cells after olaparib (10μM) treatment for 24 hours. γ-H2AX foci were counted in at least 50 cells under each condition. Three independent experiments were performed. (B) γ-H2AX, PARP and cleaved PARP protein levels were determined using Western blot in AAVS1 control, SKO and DKO cells after olaparib treatment as indicated for 24 hours. β-tubulin served as a loading control. (C) Caspase3/7 activity was measured in AAVS1 control, SKO, and DKO cells after olaparib treatment as indicated for 24 hours. (D) Representative images of immunofluorescence staining for RAD51 foci in AAVS1 control, SKO, and DKO cells after olaparib (10μM) treatment for 24 hours. RAD51 were counted in at least 50 cells for each replicate under each condition (n = 3 biological replicates). *P*-values determined by two-tailed *t* test. * *p*<0.05, ** *p*<0.01, *** *p*<0.001, **** *p*<0.0001, and not significant (ns).

We next asked whether HRR function was enhanced after RB1 loss in DKO cells. RAD51 is central to HRR, as it mediates DNA homologous pairing and strand invasion (*28*). We assessed the formation of RAD51 foci, a marker for HRR competence (*29*), by immunofluorescence staining. We observed that RAD51 foci were slightly increased after olaparib treatment for 24 hours in SKO C4-2B and 22RV1 cells as well as in their corresponding AAVS1 control cells. (Fig. 3D), indicating activation of HRR not affected by RNASEH2B deletion. However, this preserved baseline HRR function in SKO cells was not sufficient to repair DNA DSBs as we observed accumulation of γ-H2AX foci (Fig. 3A). On the other hand, we detected significantly increased RAD51 foci and less DNA DSBs in DKO cells, indicating much improved HRR capacity after RB1 loss. These results suggest that PCa cells become insensitive to PARP inhibition likely due to more efficient DNA damage repair after RB1 loss.

### Loss of RB1 upregulates HRR gene expression through E2F1 activation

We next investigated the mechanism by which HRR function was enhanced after RB1 loss. It is well-known that active form of RB1 interacts with transcription factor E2F1 and restrains its transcription activity (*30*). Loss of RB1 derepresses E2F1 activity and induces the expression of E2F1 target genes involving cell cycle progression and DNA repair (*31*). Therefore, we speculated that HRR gene expression might be upregulated through E2F1 transcriptional activation, which in turn enhanced HRR function and rendered cells resistance to PARP inhibition. In line with previous studies (*32*), we found that the transcript level of E2F1 itself was upregulated in primary and metastatic PCa tumors with RNASEH2B/RB1 co-deletion (Fig. 4A), likely due to a positive feedback loop. This was supported by the data from publicly available E2F1 chromatin immunoprecipitation sequencing (ChIP-seq) data (*33, 34*), showing strong E2F1 binding at its own promoter region (fig. S6). Using an E2F1 reporter assay, we detected significantly higher E2F1 transcriptional activity in RNASEH2B/RB1 DKO cells compared to RNASEH2B SKO cells (Fig. 4B). The E2F1 transcriptional activity remained at a high level after olaparib treatment. We further analyzed E2F1 ChIP-seq data and found strong E2F1 binding at the promoter regions of BRCA1/2 and RAD51 genes in PCa LNCaP cells (Fig. 4C). Notably, robust E2F1 ChIP-seq peaks are located immediately upstream of the transcription start sites, indicating a direct transcriptional regulation. We then performed E2F1 ChIP combined with quantitative polymerase chain reaction (qPCR) and detected enriched E2F1 binding at the promoter regions of BRCA1/2 and RAD51 genes in parental C4-2B and 22RV1 cells (Fig. 4C). To further demonstrate E2F1-mediated upregulation of BRCA1/2 and RAD51 in DKO cells, we knocked down E2F1 expression using RNA interference and observed significantly decreased BRCA1/2 and RAD51 protein levels (Fig. 4D). In addition, treatment of DKO cells with a pan-E2F inhibitor HLM006474 reduced BRCA1/2 and RAD51 protein expression. We next compared gene expression changes in RB1-deleted DKO cells relative to RB1-intact SKO. We found that the mRNA levels of BRCA1/2 and RAD51 were significantly upregulated in DKO cells (Fig. 4E). We further compared their protein levels in the absence and presence of olaparib (Fig. 4F). We detected much higher protein levels of BRCA1/2 in DKO cells, while the RAD51 protein level remained unchanged, indicating post-transcriptional regulation involved after RB1 loss in these cells. Notably, olaparib treatment suppressed BRCA1/2 expression in SKO cells, which might contribute to PARPi response in these cells. This is in agreement with the results from previous studies, showing PARP1 functions as E2F1 co-factor and regulates DNA repair gene expression (*35–37*). Nevertheless, BRCA1/2 protein levels were restored and remained at a high level after olaparib treatment in DKO cells. Considering the role of RB1-E2F1 signaling in cell cycle regulation (*34*), we performed cell cycle analysis and found a negligible change across AAVS1 control, SKO and DKO cells (Fig. 4G). Therefore, upregulation of BRCA1/2 expression is largely due to transcriptional regulation rather than cell cycle alteration after RB1 loss, although BRCA1/2 expression is cell cycle dependent. Finally, in the SU2C/PCF cohort, we observed that metastatic PCa tumors with homozygous RB1 deletions had significantly higher transcript levels of BRCA1/2 (Fig. 5A), which might have potential to repair damaged DNA more effectively and survive PARP inhibition. Taken together, our results suggest that RB1 loss upregulates BRCA1/2 gene expression through E2F1 transcriptional activation. The expression of BRCA1/2 remains at a high level after PARP inhibition, leading to proficient DNA DSB repair and PARPi resistance.

**Fig.4.**
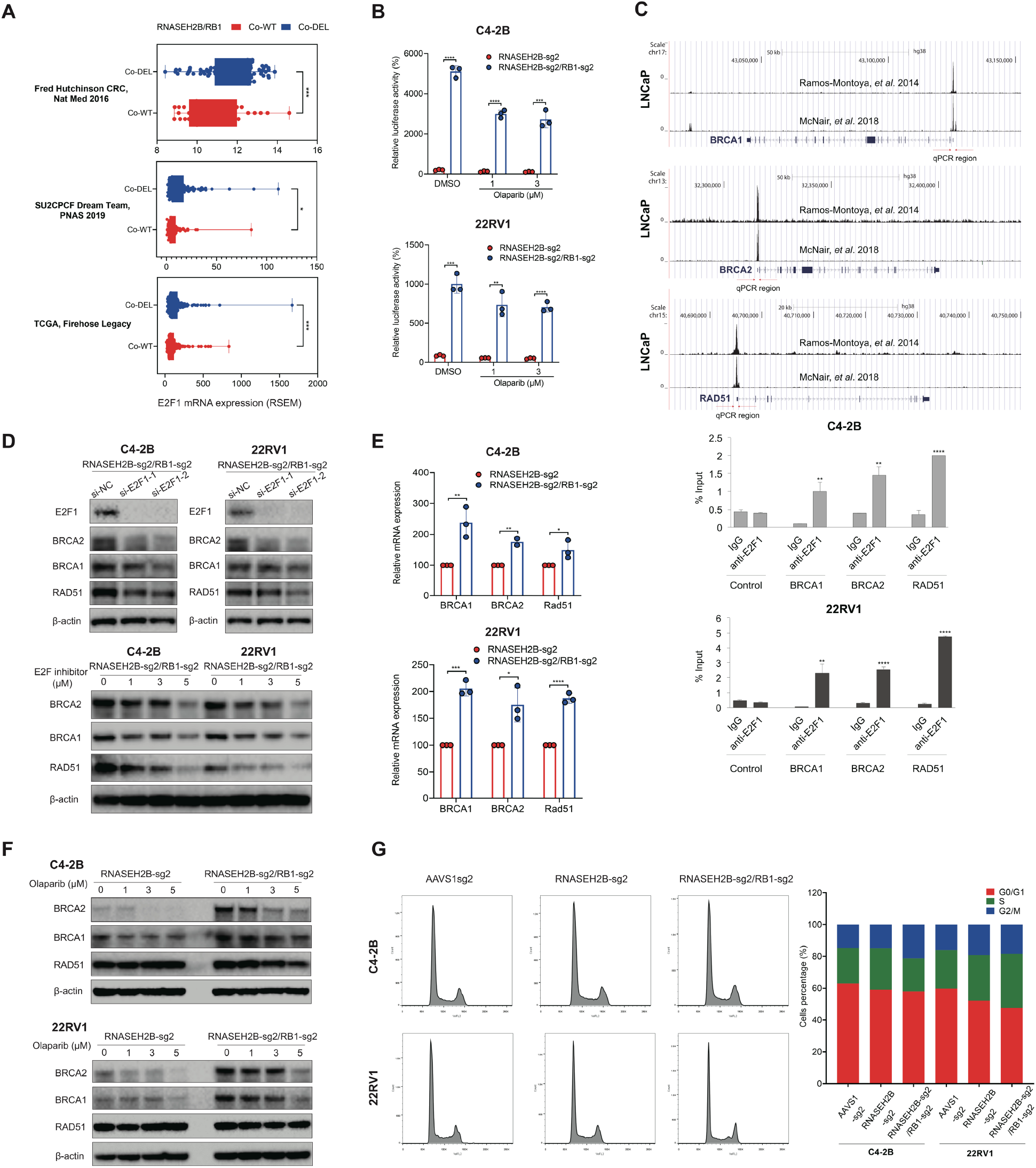
RB1 loss upregulates HRR gene expression through activating E2F1 transcriptional activity. (A) The comparison of E2F1 transcript levels between RNASEH2B/RB1 co-wildtype (co-WT) and co-deletion (co-DEL) tumors in three PCa clinical cohorts. (B) The comparison of E2F1 transcriptional activity between SKO and DKO cells in the presence or absence of olaparib as indicated for 24 hours using E2F1 luciferase reporter assay. (C) Publicly available E2F1 ChIP-seq data showing E2F1 binding capacity at the promoters of BRCA1/2 and RAD51 genes in PCa LNCaP cells. E2F1 binding was determined by ChIP-qPCR at the promoters of BRCA1/2 and RAD51 genes in C4-2B and 22RV1 cells. Red arrows indicate the qPCR regions. Normal IgG was used as a control in contrast to anti-E2F1 antibody. (D) Western blot showing protein levels of indicated genes in DKO C4-2B and 22RV1 cells after E2F1 siRNA knockdown or the treatment with pan-E2F inhibitor HLM006474 as indicated for 24 hours. β-actin served as a loading control. (E) BRCA1/2 and RAD51 mRNA levels were determined by RT-qPCR in DKO relative to SKO C4-2B and 22RV1 cells. (F) Western blot showing protein levels of BRCA1/2 and RAD51 in SKO vs. DKO cells after olaparib treatment for 24 hours. (G) Cell cycle distribution in AAVS1 control, SKO and DKO C4-2B and 22RV1 cells under regular cell culture condition. *P*-values determined by two-tailed *t* test. * *p*<0.05, ** *p*<0.01, *** *p*<0.001, and **** *p*<0.0001.

**Fig.5.**
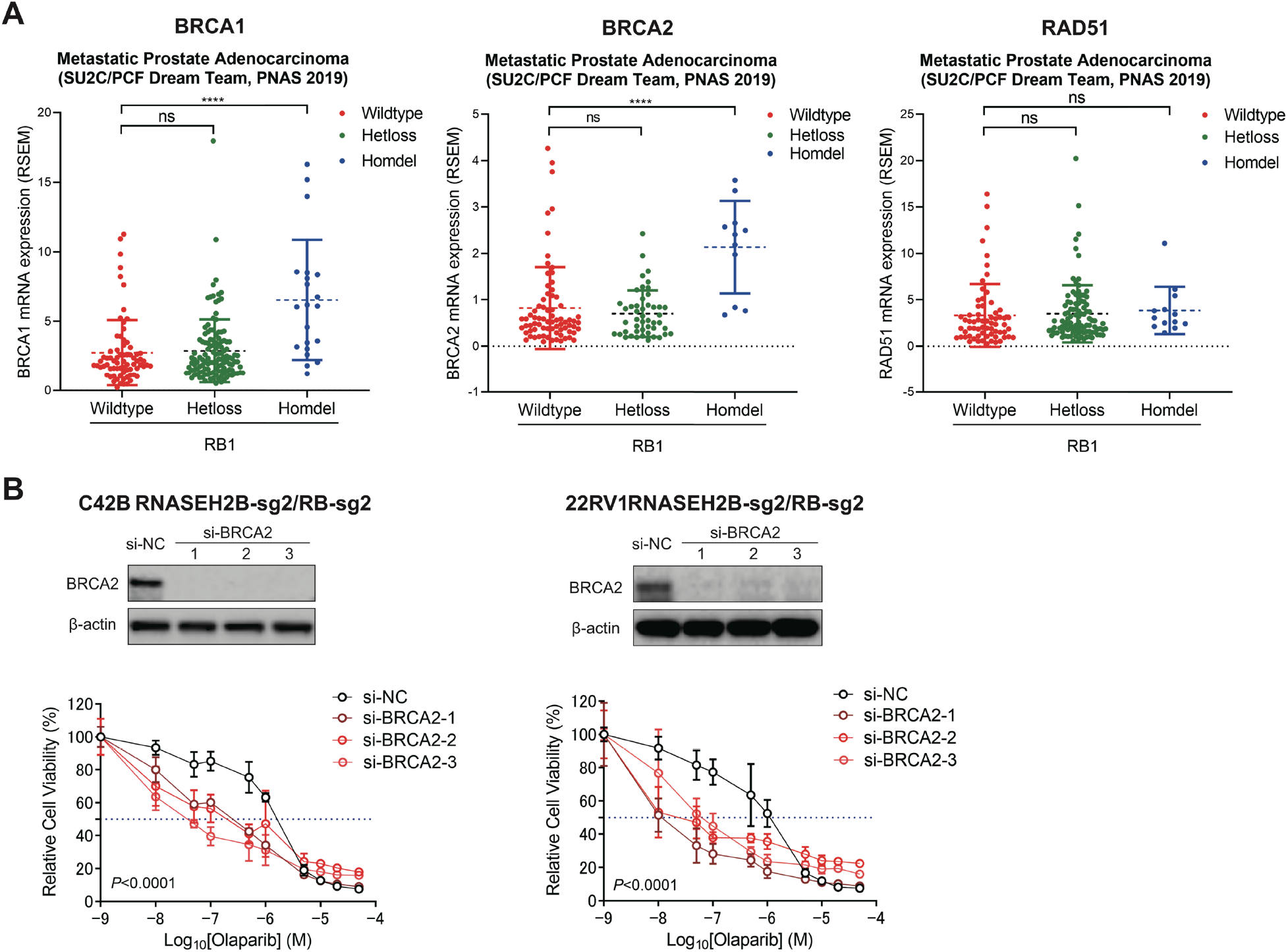
Loss of BRCA2 re-sensitizes RNASEH2B/RB1 DKO cell to PARP inhibition. (A) mRNA expression levels of BRCA1/2 and RAD51 in metastatic PCa tumors (in the SU2C/PCF cohort) harboring wildtype RB1, heterozygous (Hetloss), and homozygous (Homdel) RB1 deletions. Tumors with BRCA1/2 and RAD51 deletions were excluded in each analysis respectively. *P*-values determined by two-tailed *t* test **** *p*<0.0001 and not significant (ns). (B) RNASEH2B/RB1 DKO C4-2B and 22RV1 cells were transfected with three different BRCA2 siRNA or a negative control (NC) siRNA at final concentration of 10 nM for 2 days, followed by the treatment with the indicated doses of olaparib for additional 7 days. Cell viability was determined using alamarBlue assay (mean ± SD; n=3). Western blot showing BRCA2 protein levels 48 hours after siRNA transfection. *P*-values determined by two-way ANOVA.

### PCa cells with co-loss of RNASEH2B/RB1/BRCA2 are sensitive to PARP inhibition

While BRCA1 is critical in HRR, genomic alterations in PCa involve BRCA2 more commonly than BRCA1. Clinical next-generation sequencing analyses of both primary and metastatic PCa tumors have revealed that BRCA2 is co-deleted with RNASEH2B and RB1 in a small portion of patients (Fig. 1D). Importantly, our data have suggested that upregulation of BRCA2 through the RB1-E2F1 pathway likely contributes to PARPi resistance in RB1-deleted cells. We next asked whether deletion of BRCA2 can re-sensitize DKO cells to PARP inhibition. Here, we knocked down BRCA2 expression in RNASEH2B/RB1 DKO C4-2B and 22RV1 cells using RNA interference. Three different siRNAs against BRCA2 completely abolished BRCA2 protein expression determined by Western blot (Fig. 5B). We found that depletion of BRCA2 renders DKO C4-2B and 22RV1 cells sensitivity to olaparib, indicating that elevated BRCA2 expression after RB1 loss is likely one of the mechanisms for PARPi resistance. Importantly, RNASEH2B/RB1 DKO cells also respond to other PARPis (veliparib, rucaparib, and talazoparib) following BRCA2 depletion (fig. S7). These results suggest that BRCA2-deficient tumors may respond to PARPis regardless RB1 status.

### ATR inhibition overcomes PARPi resistance of PCa tumors with RNASEH2B/RB1 co-deletion

Since PCa cells with RNASEH2B single gene deletion or RNASEH2B/RB1/BRCA2 triple gene deletion are hypersensitive to PARP inhibition, we next asked how to overcome PARPi resistance for cells with RNAEH2B/RB1 double gene deletion. Patients with tumors harboring RNASEH2B/RB1 co-deletion account for 10.6% and 3.2% in all primary and metastatic PCa cases, respectively (Figure 1D). Emerging evidence has shown that PARP inhibition may activate ATR, which phosphorylates and activates CHK1 and allows cells to survive PARPi-induced replication stress (*38*). Previous studies have also demonstrated that ATR-CHK1 signaling controls E2F-dependent transcription of HRR genes (*39–41*). Here, we found that ATR activity was elevated in RNASEH2B/RB1 DKO cells after olaparib treatment, as evidenced by increased CHK1 phosphorylation in a dose-dependent manner (Fig. 6A). We hypothesized that DKO cells relied on ATR activity to survive PARPi-induced DNA damage. We therefore sought to investigate the effect of PARPi and ATR inhibitor (ATRi) either alone or in combination on the growth of DKO cells. To achieve ATR inhibition, we utilized a clinically used ATR inhibitor (ATRi) VE-822. Both SKO and DKO cells fail to show increased response to VE-822 as a single agent in comparison to AAVS1 control cells (Fig. 6B). We treated PARPi-insensitive DKO cells with olaparib combined with VE-222 and found co-treatment diminished the growth of these cells (Fig. 6C). We observed that DKO C4-2B and 22RV1 cells were re-sensitized to olaparib in the context of ATR inhibition (Fig. 6D). Using the Loewe and Bliss Synergy analysis (*42, 43*), we found a synergistic interaction between olaparib and VE-822, with a high synergy score for DKO 22RV1 (Loewe: 13.263; Bliss: 16.347) and C42B (Loewe: 8.314; Bliss: 13.502) cells (Fig. 6E). Synergistic effects were also observed in the same cells using colony formation assay (Fig. 6F).

**Fig.6.**
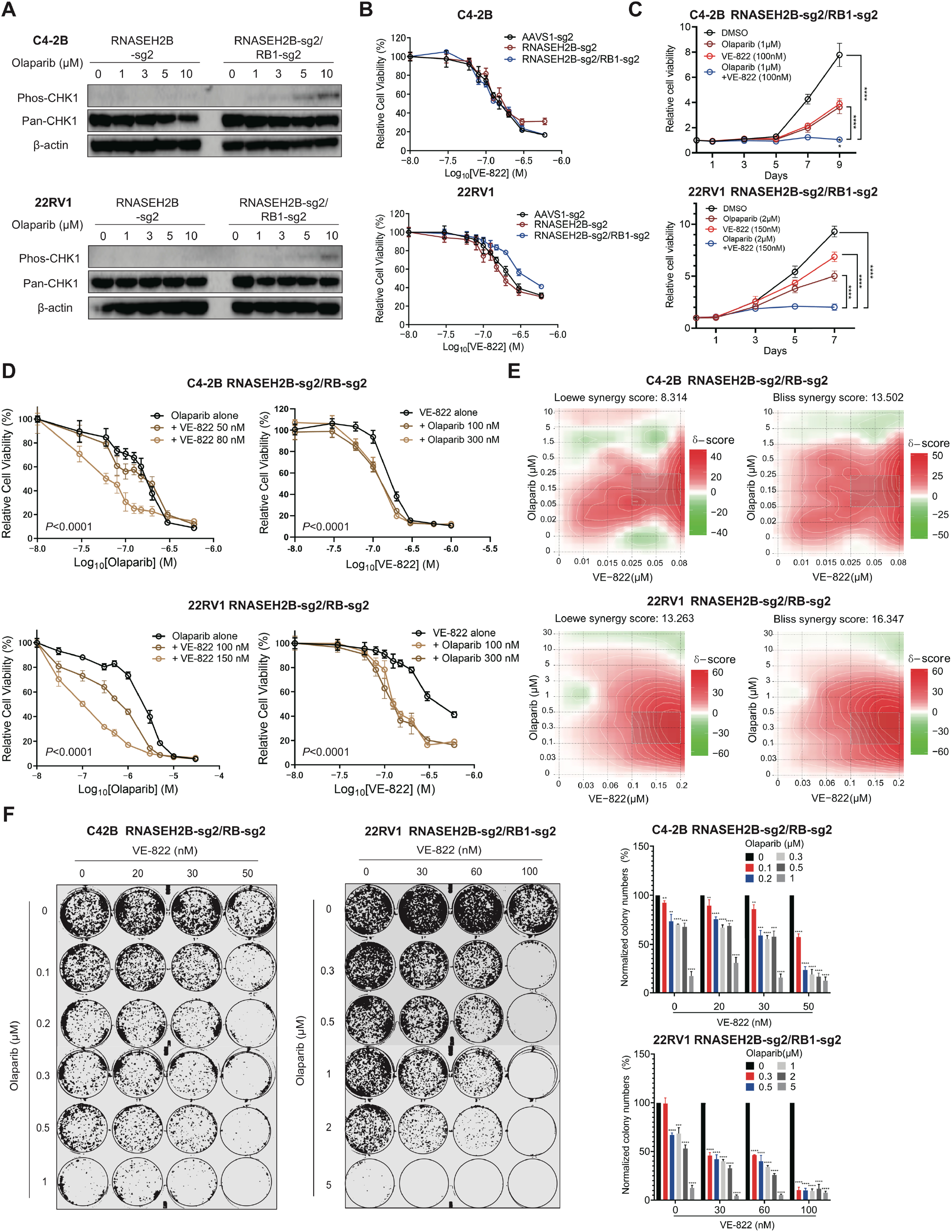
ATR inhibition overcomes PARPi resistance in RNASEH2B/RB1 DKO cells. (A) Western blot showing phosphorylated CHK1 and total CHK1 protein levels in RNASEH2B SKO and RNASEH2B/RB1 DKO cells after the treatment with the indicated doses of Olaparib for 24 hours. (B) Cell viability of AAVS1 control, SKO and DKO cells was determined using alamarBlue assay after the treatment with ATR inhibitor VE-822 as indicated for 7 days. (C) RNASEH2B/RB1 C4-2B and 22RV1 DKO cells were treated with olaparib, VE-822, Olaparib + VE-822 as indicated. Cell proliferation rates were determined using alamarBlue assay. (D) RNASEH2B/RB1 DKO cells were treated with olaparib alone or in combination with VE-822 as indicated for 7 days. Cell viability was determined using alamarBlue assay. (E) The synergistic score between olaparib and VE-822 was determined using Loewe and Bliss Synergy analysis. (F) The growth of RNASH2B/RB1 DKO cells after the treatment with olaparib and/or VE-822 for 14 days was determined using colony formation assay. Colony number was quantified using ImageJ. *P*-values determined by two-tailed *t* test. ** *p*<0.01, *** *p*<0.001, and **** *p*<0.0001.

Next, we tested combination treatment *in vivo* using PARPi-insensitive DKO 22RV1 cells. After xenograft tumors established in ICR-SCID mice, animals were divided into four group and treated with vehicle, olaparib, VE-822, or olaparib in combination with VE-822 for 3 cycles as indicated (Fig. 7A). We found that tumor growth was significantly inhibited by combination treatment, while both olaparib and VE-822 had little effect as a single agent. No significant mouse weight loss was observed in all four groups, indicating the combination treatment is tolerable.

**Fig.7.**
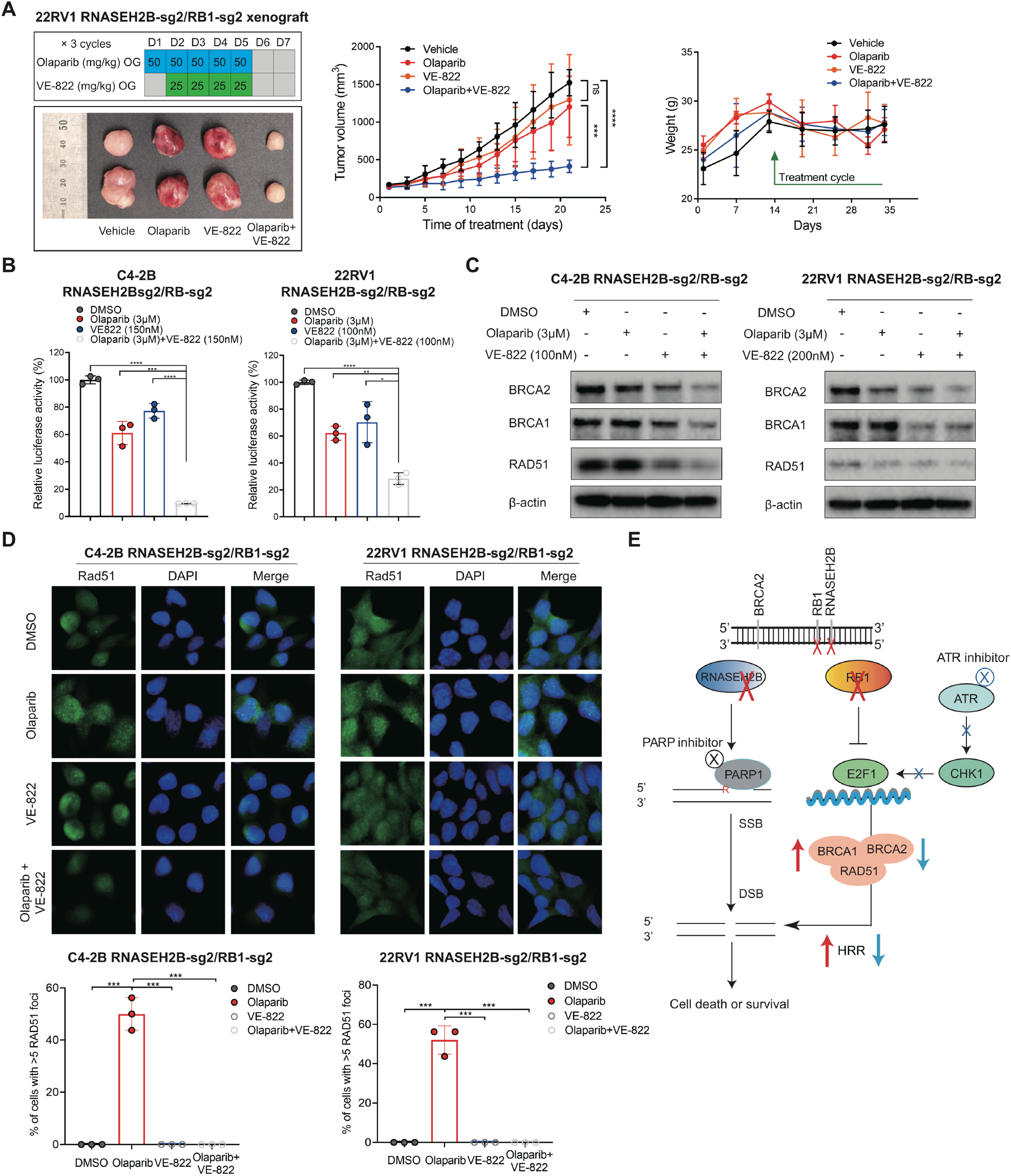
Combination therapy with ATR and PARP inhibition suppresses RNASEH2B/RB1 DKO cell growth *in vivo*. (A) RNASEH2B/RB1 DKO 22RV1 cells were injected subcutaneously into ICR-SCID mice. Mice were randomly assigned into four groups (n=5 animals per group) and treated with vehicle, olaparib (50mg/kg), VE-822 (25mg/kg), or olaparib in combination with VE-822 for 3 cycles as indicated. Both drugs were administered by oral gavage (OG) once a day. Tumor volume and mouse weight were recorded and analyzed across four groups as indicated. (B) DKO PCa cells were treated with DMSO, Olaparib, VE-822 or Olaparib + VE-822 for 24 hours. E2F1 activity was detected using E2F1 luciferase reporter assay. (C) Western blot showing protein levels of BRCA1/2 and RAD51 in DKO cells after the treatment with DMSO, Olaparib, VE-822 or Olaparib + VE-822 for 24 hours. (D) Representative images of immunofluorescence staining for RAD51 foci in DKO C4-2B and 22RV1 cells after the treatment with DMSO, olaparib (10μM), VE-822 (1000nM) or Olaparib + VE-822 for 24 hours. olaparib (10μM) treatment for 24 hours. RAD51 were counted in at least 50 cells for each replicate under each condition (n = 3 biological replicates). (E) Schematic model depicting the mechanism by which concurrent deletions of RNASEH2B, RB1, and BRCA2 genes impact the response to PARP inhibition. *P*-values determined by two-tailed *t* test. * *p*<0.05, ** *p*<0.01, *** *p*<0.001, **** *p*<0.0001, and not significant (ns).

We then asked whether the combination of PARP and ATR inhibition affects E2F1-mediated BRCA1/2 expression and HRR function. Using an E2F1 reporter assay, we observed significantly decreased E2F1 activity in DKO C4-2B and 22RV1 cells after the combination treatment (Fig. 7B). The protein expression levels of BRCA1/2 and RAD51 were also decreased after combination treatment (Fig. 7C). Finally, we examine HRR function using RAD51 foci formation assay in DKO cells. We observed increased RAD51 foci after olaparib treatment, which was diminished by combined treatment with VE-822 (Fig. 7D). The loss of RAD51 foci after ATR inhibition is likely due to reduced BRCA1/2 expression (Figure 7C) and disrupted BRCA-independent RAD51 loading to DSBs as previously reported (*44*). Taken together, our results support the notion that the combined therapy with PARP and ATR inhibitors may overcome PARPi resistant in PCa cells with RNASEH2B/RB1 co-deletion through inhibition of E2F1-mediated augmentation of HRR capacity.

## Discussion

It has been a great challenge to determine which patients are most likely to benefit from PARP inhibition. Preclinical studies and clinical trials have demonstrated that PCa patients with tumors harboring deleterious germline or somatic BRCA1/2 alterations have a high response rate to PARPis. On the other hand, alterations in other HRR genes [known as BRCAness genes (*45*)], such as ATM and CHEK2, are not associated with response to the same extent. Furthermore, PARPi response for tumors harboring genomic alterations in non-HRR DDR genes remains largely unknown. RNASEH2B is not a BRCAness gene. Instead, it is one of three genes encoding RNase H2 protein complex, which cleaves single ribonucleotides incorporated into the genome. In this study, we show that RNASEH2B is frequently deleted in both primary and metastatic PCa tumors, which creates DNA lesions and increases PARP trapping, leading to accumulation of DNA DSBs and apoptotic cell death after the treatment with PARPis. While RNASEH2B deletion is an attractive biomarker to predict PARPi response in PCa, co-deletion with RB1 counteracts the cytotoxic effects of PARP trapping, at least in part, by upregulation of E2F1-mediated BRCA1/2 expression, thereby enhancing HRR capacity (Fig. 7E). Subsequently, we show that deletion of BRCA2 re-sensitizes RNASEH2B/RB1 co-deleted cells. Our work further demonstrates that the combination of PARP and ATR inhibition can overcome intrinsic PARPi resistance rising from RB1 loss. Taken together, these results provide a basis of clinical application of PARPi either alone or in combination with ATRi, given the interaction between multiple genomic alterations in PCa. Patients will likely respond to PARP inhibition if their tumors harbor RNASEH2B singe deletion or RNASEH2B/RB1/BRCA2 triple co-deletion, whereas patients with tumors harboring RNASEH2B/RB1 double co-deletion may respond to combined PARP and ATR inhibition.

Loss of RB1 has been shown to be strongly associated with poor clinical outcomes in advanced PCa by facilitating lineage plasticity in the context of concurrent loss of TP53 (*46–49*). RB1/TP53-deficient tumors are resistant to a wild range of single agent therapeutics, including PARPi talazoparib (*32*). Furthermore, studies have shown that the combination of PARP inhibition and RB1-associated CDK inhibition may be a viable strategy for neuroendocrine PCa treatment (*50*), supporting an important role of RB1 in PARP inhibition. Our present work does not exclude the possibility that PARPi resistance after RB1 loss results from lineage plasticity and epigenetic reprogramming (*51*). Nevertheless, tumors with RB1 loss express significantly higher levels of E2F1, which directly regulates HRR genes, most notably BRCA1/2. Our results strongly suggest that RB1 loss renders PARP inhibition inefficient for tumors with non-BRCA genomic alterations, because E2F1-induced BRCA1/2 expression enhances HRR capacity. Genomic aberrations of RB1 are common as late, subclonal events in mCRPC (*52*). This may partially explain why mCRPC patients with tumors harboring alterations in non-BRCA HRR genes have lower response rate. On the other hand, tumors with BRCA1/2 alterations remain sensitive to PARP inhibition regardless RB1 status (*53*). These tumors appear unaffected by changes in cell context or genetic background due to lack of buffering mechanisms.

The landscapes of cancer gnome are complex including base changes, indels, copy number changes, and structural rearrangements. Genomic deletion is common in cancer and ranges from focal deletions affecting a few genes to arm-level deletions affecting hundreds to thousands of genes (*54*). Little is known about the functional consequences of large-scale genomic deletions, and it is difficult to determine the specific genes responsible for the biological effects. One of the limitations in our studies is that we didn’t test whether co-deletions of other protein-coding genes, let alone non-coding RNAs, on chromosome 13q may also impact PARPi sensitivity. While CRISPR screens have not identified any proximal genes at the RNASEH2B/RB1/BRCA2 loci, which when deleted cause increased sensitivity or resistance to PARP inhibition, further investigation is needed by creating isogenic cell lines with engineered large-scale deletions instead of a gene-by-gene approach. Accordingly, there is a rationale for examining copy number changes and structural rearrangements of PCa tumors, which are not captured by targeted next-generation sequencing tests being implemented in current clinical practice.

The mechanisms of acquired PARPi resistance has been heavily studied in BRCA-deficient cells. A key mechanism appears to be the restoration or bypass of HRR and fork protection functions, which can be overcome by ATR inhibition (*44*). In our studies, we propose an intrinsic resistant mechanism through the RB1-E2F1-BRCA pathway in non-BRCA deficient cells. We demonstrate that ATR inhibition may impair E2F1-induced BRCA1/2 expression and re-sensitize cells to PARPis. The combination therapy with PARPi and ATRi is being evaluated in clinical trials for mCRPC patients (NCT03787680). Therefore, it is conceivable to develop predictive biomarkers based on a comprehensive genomic testing and explore the combination of PARP and ATR inhibition as a promising strategy for advanced PCa patients when a single agent fails.

## Materials and Methods

### Cell lines and materials

Human PCa cell lines LNCaP, C4-2B, 22Rv1, PC-3, and DU145 (American Type Culture Collection, ATCC) were cultured in RPMI1640 medium (Thermo Fisher Scientific), while 293FT cells (Thermo Fisher Scientific) were maintained in DMEM medium (Thermo Fisher Scientific). Media were supplemented with 10% fetal bovine serum (Sigma-Aldrich), 1% penicillin/streptomycin (Sigma-Aldrich), and 1% HEPES (Sigma-Aldrich). All cell lines were authenticated using high-resolution small tandem repeats (STRs) profiling at Dana-Farber Cancer Institute (DFCI) Molecular Diagnostics Core Laboratory and were tested mycoplasma-free before experiments. The small molecule inhibitors are listed in table S3.

### Establishment of CRISPR/Cas9 KO cell lines

CRISPR guides targeting RNASEH2B were cloned into lentiGuide-Puro vector (#52963, Addgene), while CRISPR guides targeting RB1 were cloned into lenti-sgRNA hygro vector (#104991, Addgene). The lentiCas9-Blast vector that expresses Cas9 was obtained from Addgene (#52962). Lentiviruses were generated using packaging vectors pMD2.G (#12259, Addgene) and psPAX2 (#12260, Addgene) with Lipofectamine^TM^3000 transfection reagents (#L3000015, Invitrogen) in 293FT cells. PCa cells were initially infected with lentiviruses of Cas9 and selected with Blasticidin (10 μg/ml) for two weeks. Polybrene was added at a final concentration of 8ug/ml to increase transduction efficiency. To generate RNASEH2B-KO cells, PCa cells were infected with lentiviruses containing specific sgRNAs and selected with puromycin (3μg/ml) for two weeks. The RNASEH2B-KO cells were infected with lentiviruses with RB1 sgRNAs to generate RNASEH2B/RB1 co-deletion cells. Cells were further selected using hygromycin (300 μg/ml) for 2 weeks. sgRNA sequences are listed in table S3.

### Cell viability assays

PCa cells were seeded in 96-well plates (1-2 ×10^3^cells/well) and treated as indicated. Cell viability was measured using alamarBlue Cell Viability Reagent (DAL1100, Thermo Fisher Scientific) according to the manufacturer’s instructions.

### Colony formation assays

PCa cells were seeded in 12-well plates at low density to avoid contact between clones (3000 cells per well). Subsequently, cells were treated as indicated 18 hours after attachment and allowed to grow for 14 days. Colonies were fixed with paraformaldehyde (4%) for 10 minutes and stained with crystal violet (1%) for 15 minutes. Colony images were quantified using ImageJ software (National Institutes of Health).

### Western blot assays

PCa cells were treated as indicated and harvested for protein extraction. Cells were rinsed with phosphate-buffered saline (PBS), scaped and lysed in cold RIRA Lysis and Extraction Buffer (#89900, Thermo Fisher Scientific) containing Protease and Phosphatase Inhibitor Cocktail (#78447, Thermo Fisher Scientific). Protein concentration was quantified using Pierce BCA Protein Assay Kit (#23225, Thermo Fisher Scientific) and measured with a spectrophotometer. Western blot was performed as previously described (*55*). Antibodies were listed in table S3.

### ChIP-qPCR assays

ChIP experiments were performed as previously described (*55, 56*). Briefly, PCa cells were grown in RPMI1640 medium with 10% fetal bovine serum for 2 days prior to ChIP. Cells were cross-linked by formaldehyde (1%) at room temperature (RT) for 10 minutes. After washing with ice- cold PBS, cells were collected and lysed. The soluble chromatin was purified and fragmented by sonication. Immunoprecipitation (IP) was performed using normal IgG or E2F1 antibody (2 μg/IP). ChIP DNA was extracted and analyzed by qPCR using iTaq Universal SYBR Green Supermix (#1725120, Bio-Rad). The antibodies and primer sequences are listed in Table S3.

### RT-qPCR assays

Total RNA was extracted using Trizol Reagent (#15596026, Thermo Fisher Scientific) according to the manufacturer’s protocol. RT-qPCR assay were performed as previously described (*57*). Primer sequences are listed in table S3.

### Caspase-3/7 activity assays

PCa cells were seeded in 96-well plates and treated with DMSO or specific inhibitors as indicated. Caspase-3/7 activity was measured using Caspase-Glo 3/7 Assay Systems (G8091, Promega) according to the manufacturer’s protocol.

### E2F1 reporter activity assays

The E2F luciferase reporter plasmid was described previously (*55*). The reporter construct contains three tandem E2F1 consensus elements — TGCAATTTCGCGCCAAACTTG — (*58*), subcloned into SacI/XhoI sites of the pGL4.26 vector (Promega) upstream of a minimal promoter. Cells (1 x10^4^ cells/well) were seeded into 96-well plates and transfected with E2F1 luciferase reporter plasmids (50 ng/well) using X-tremeGENE HP DNA Transfection Reagent (#06366236001, Sigma-Aldrich). After 12 hours, cells were treated with DMSO or specific inhibitors as indicated for additional 24 hours. The luciferase activity was measured using One-Glo Luciferase Assay System (E6110, Promega) according to the manufacturer’s protocol..

### RNA interference

PCa cells were transfected with siRNAs (at a final concentration of 10 nM) using Lipofectamine RNAiMAX Transfection Reagent (#13778150, Thermo Fisher Scientific) according to the manufacturer’s instructions. All siRNAs were purchased from Sigma-Aldrich and listed in table S3. Cell viability and Western blot assays were performed 2 days after siRNA transfection.

### Immunofluorescence staining

PCa cells were seeded onto the Millicell EZ SLIDE 4-well glass slides (PEZGS0496, Millipore) pre-coated with Poly-L-lysine (P4707, Sigma-Aldrich) and then processed with treatments as indicated. After 24 hours, cells were washed with PBS and fixed in 4% formaldehyde at RT for 10 minutes. Fixed cells were washed with PBS for 3 times and extracted with 0.2% Triton X-100 for 10 minutes. Subsequently, cells were blocked in blocking buffer (5% bovine serum albumin in PBS) for 1 hour and incubated with the primary antibody RAD51 (ab133534, Abcam) or phosphor-H2AX (#05-636, Sigma-Aldrich) at 1:200 dilution. Following an overnight incubation at 4°C, slides were washed with PBS and incubated with Alexa Fluor 488-conjugated anti-mouse (A28175, Life Technologies) or anti-rabbit (A27034, Life Technologies) secondary antibody at 1:1000 dilution at RT for 1 hour. After washing, the Mounting Medium with DAPI (ab104139, Abcam) was applied onto the slides. The slides were imaged under fluorescence microscope and quantified using ImageJ software. γ-H2AX was quantified by counting the number of foci per cell, while RAD51 was quantified by scoring the percentage of cells with ≥ 5 foci/cell. Each experiment was performed in triplicate independently, and at least 50 cells were counted for each replicate under each condition.

### Xenograft experiments

RNASEH2B/RB1 DKO 22RV1 cells were used to generate xenograft tumors. Cells (2.5 × 10^6^ cells/50μL/mouse with additional 50μL Matrigel) were subcutaneously injected into the right flank of male ICR-SCID mice (Taconic Laboratories) at the age of 4-5 weeks. All procedures were performed in compliance with the guidelines from the Institutional Animal Care and Use Committee (IACUC) at the Brigham and Women’s Hospital. The tumor growth and mouse body weight were monitored twice a week. Tumor volume was measured using a vernier caliper and calculated according to the formula: volume = 1/2(length×width^2^). Mice bearing about 150 mm^3^ tumors were randomized into four groups and treated with vehicle, olaparib (50 mg/kg), VE-822 (25 mg/kg), or the combination of olaparib and VE-822. Olaparib was formulated in 5% dimethylacetamide/10% Solutol HS 15/85% PBS; VE-822 was formulated in 10% Vitamin E d-alpha tocopheryl polyethylene glycol 1000 succinate (TPGS). Both drugs were administered by oral gavage once a day, with olaparib five days on-two days off and VE-822 four consecutive days a week starting at the next day of olaparib treatment. Animals were euthanized after 3-week drug treatment, or when tumors exceeded 1000 mm^3^.

### Statistical analyses

Quantitative measurements are presented as mean ± standard deviation (SD) from at least three biological replicates unless stated otherwise. Statistical analyses were performed using an unpaired two-tailed Student’s *t* test or a two-way ANOVA with a post-hoc Tukey’s honest significant difference (HSD) test when comparing at least three groups. P-values of less than 0.05 were considered as statistically significant.

### Clinical cohort analyses

Bioinformatic analysis of genomic deletion, mRNA expression and copy number variations of related genes were performed using publicly available clinical datasets in cBioPortal (*24, 25*). Statistical analyses were performed using an unpaired two-tailed Student’s *t* test.

## Acknowledgments

We thank B. Gui, C. Feng, X. Bai, T. Tian, K. Jia, and J. Geng for insightful discussion and technical support.

## Funding

National Institutes of Health grant 1R21CA252578-01 (to L.J.) National Institutes of Health grant 1R01CA262524-01 (to L.J.)

## Author contribution

L. J. conceived the project, designed the experiments, analyzed the data, and wrote the paper. C.M., T.T. performed the experiments, analyzed the data, and wrote the paper. T.T., F.G., and T.T. performed the experiments. Z.S., Z.S., K.W.M., L.Z., and A.S.K. analyzed the data and edited the manuscript.

## Competing interests

Authors declare that they have no competing interests.

## Data and materials availability

All data needed to evaluate the conclusions in the paper are present in the main text or the Supplementary Materials. Additional data related to this paper may be requested from the authors.

## Supplementary Materials

Supplementary Material for this article is available.

## Notes

### Competing Interest Statement

The authors have declared no competing interest.

