## Supplementary Figures S1-S7, Table S1-S3 for "RB1 loss overrides PARP inhibitor sensitivity driven by RNASEH2B loss in prostate cancer"

#### This PDF file include:

Figs. S1 to S7
Tables S1 to S3

**Figure S1.**

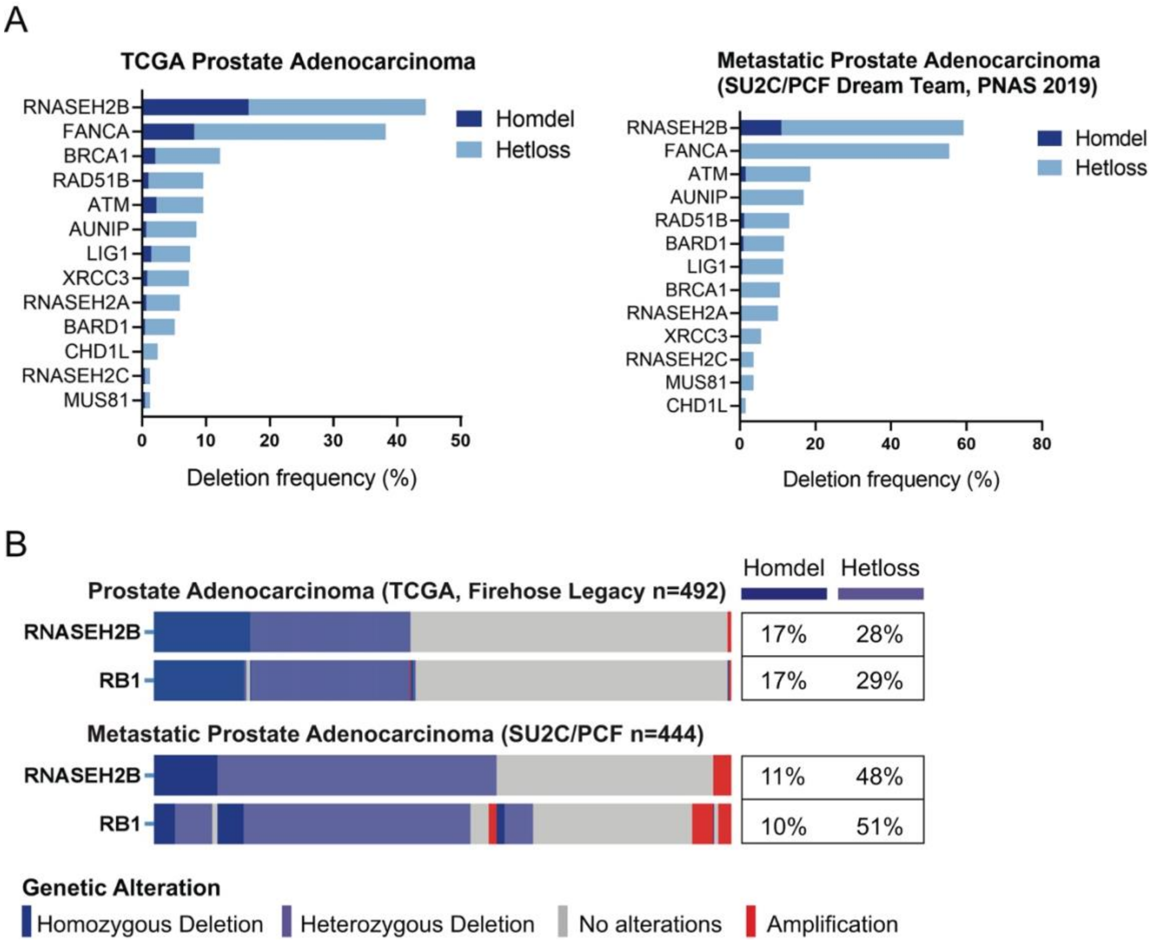


**Fig. S1. The frequency of homozygous and heterozygous deletions in genes identified from CRISPR/Cas9 screens.** (A) The frequency of homozygous (Homdel) and heterozygous (Hetloss) deletions in 13 common genes identified from CRISPR/Cas9 screens in the TCGA and SU2C/PCF cohorts. (B) the frequency of RNASEH2B and RB1 homozygous and heterozygous deletions in the TCGA and SU2C/PCF cohorts.

**Figure S2.**

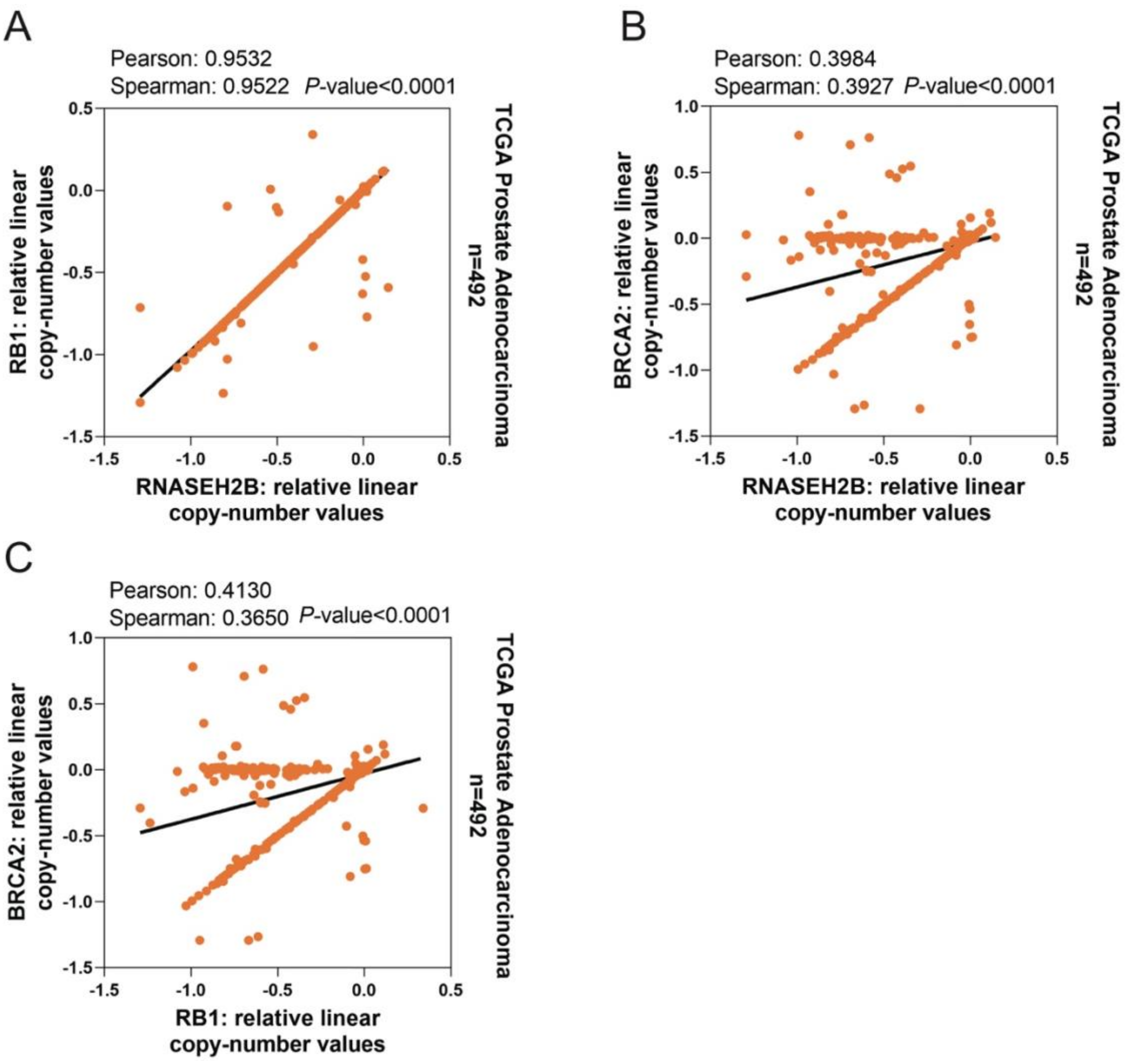


**Fig. S2. Correlation of copy numbers between RNASEH2B, RB1, and BRCA2 genes.** (A) Correlation of copy numbers between RNASEH2B and RB1 genes in the TCGA cohort. (B) Correlation of copy numbers between RNASEH2B and BRCA2 genes in the TCGA cohort. (C) Correlation of copy numbers between RB1 and BRCA2 genes in the TCGA cohort. All data were obtained from cBioPortal.

**Figure S3**

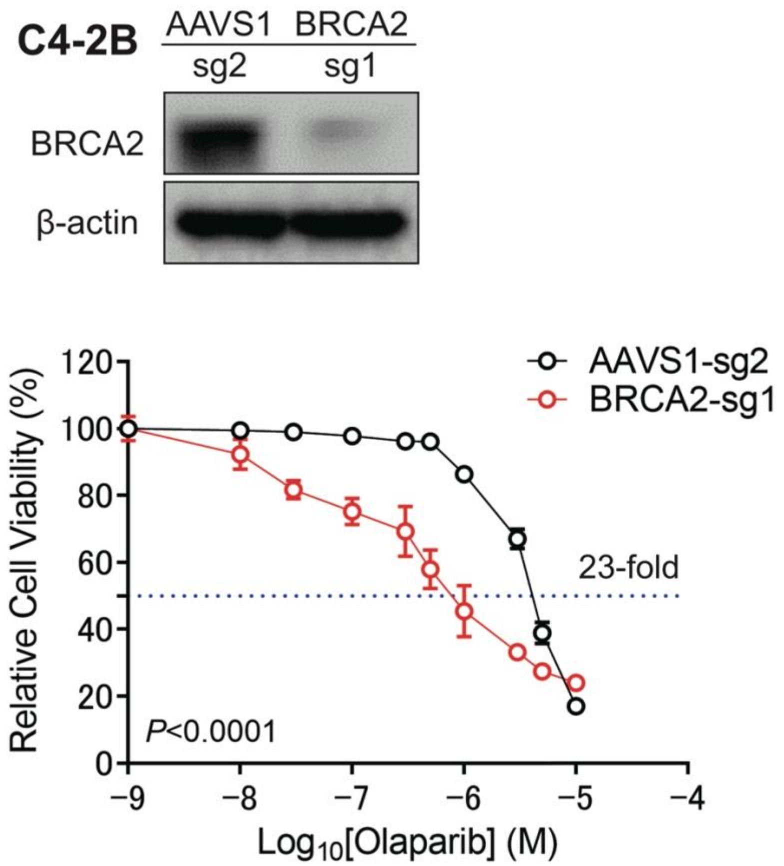


**Fig. S3. BRCA2 deletion increases C4-2B cell sensitivity to olaparib.** The BRCA2 gene was deleted in C4-2B cells using CRISPR/Cas9 gene editing. The knockout (KO) efficiency was determined by Western blot. BRCA2-KO and corresponding AAVS1 control C4-2B cells were treated with olaparib as indicated for 7 days. Cell viability was measured using alamarBlue assay. Olaparib sensitivity (determined by IC50) was increased 23-fold after BRCA2 deletion. *P*-value was determined by two-way ANOVA.

**Figure S4.**

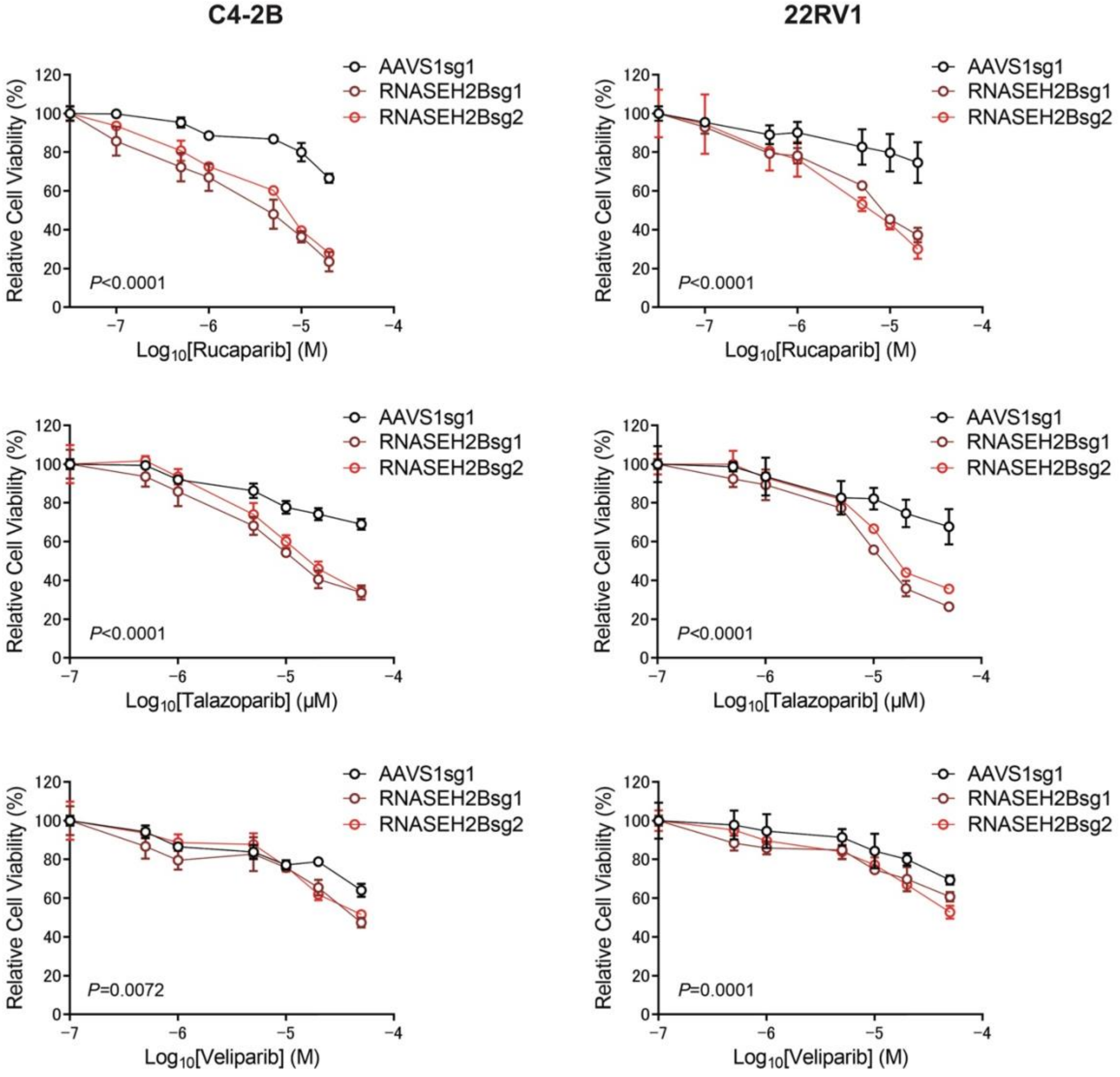


**Fig. S4. RNASEH2B-KO cells respond to PARP inhibitors with trapping ability**. RNASEH2B-KO and corresponding AAVS1 control C4-2B and 22RV1 cells were treated with talazoparib, rucaparib and veliparib as indicated for 7 days. Cell viability was measured using the alamarBlue assay kit. Two different single guide RNAs (sg1 and sg2) were used to generate RNASEH2B-KO cells. *P*-values were determined by two-way ANOVA.

**Figure S5.**

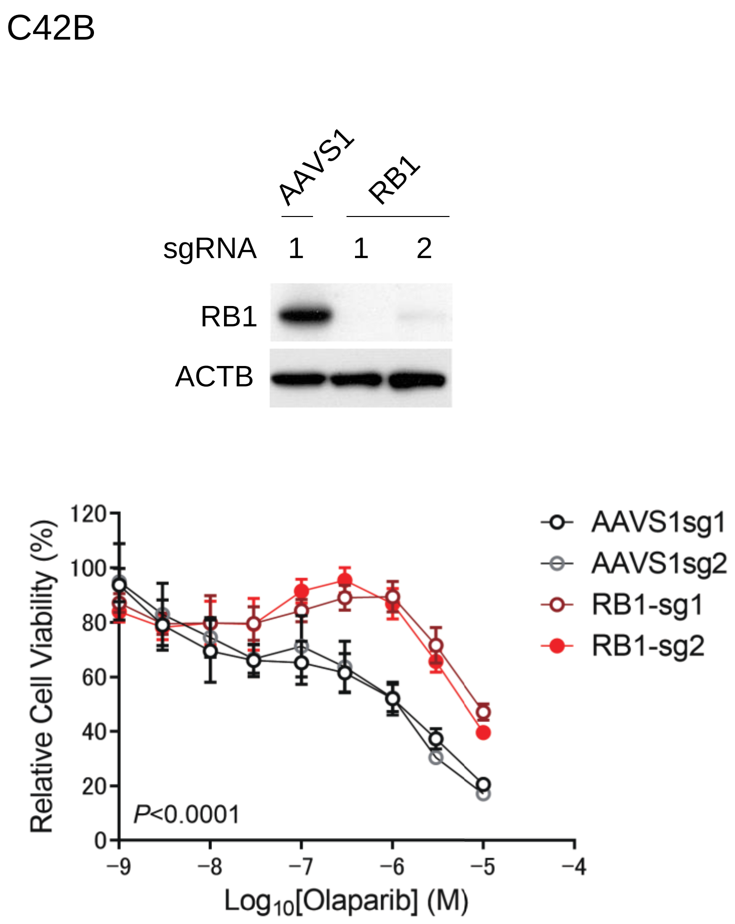


**Fig. S5. RB1 deletion reduces C4-2B cell sensitivity to olaparib.** The RB1 gene was deleted in C4-2B cells using CRISPR/Cas9 gene editing. Two different single guide RNAs (sg1 and sg2) were used to generate the KO cells. The KO efficiency was determined by Western blot. RB1-KO and corresponding AAVS1 control C4-2B cells were treated with olaparib as indicated for 7 days. Cell viability was measured using alamarBlue assay. *P*-value was determined by two-way ANOVA.

#### Figure S6.

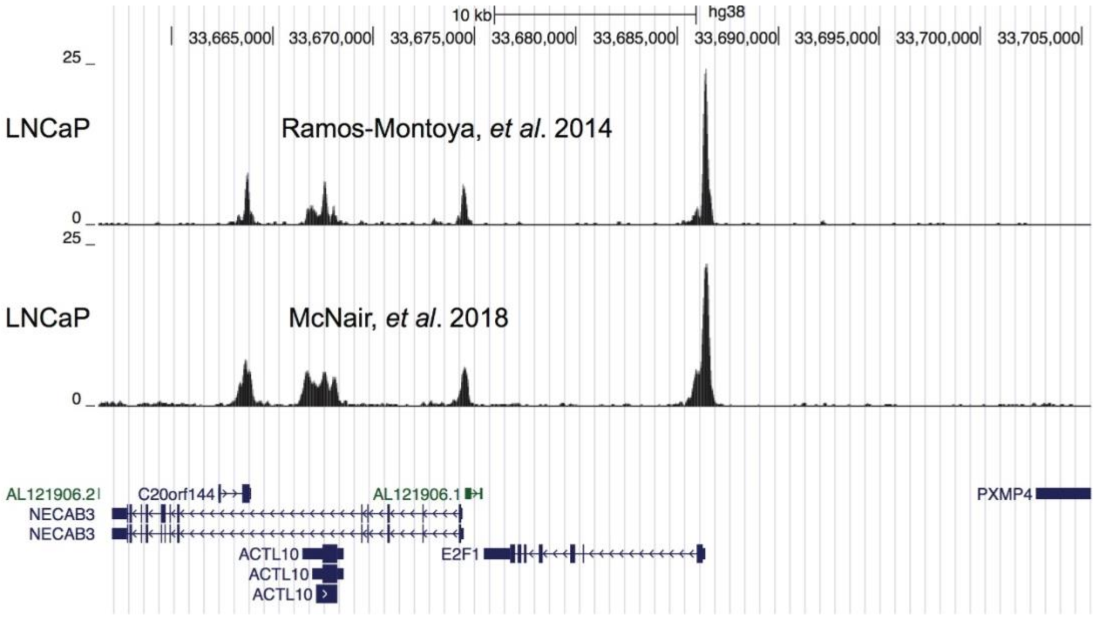


**Fig. S6. E2F1 transcription factor binds the E2F1 promoter in ChIP-seq.** Two publicly available E2F1 ChIP-seq datasets using LNCaP cells were analyzed (References *33, 34*). The E2F1 ChIP-seq peaks were observed in the UCSC Genome Browser, showing E2F1 binding capacity at the E2F1 promoter region.

**Figure S7.**

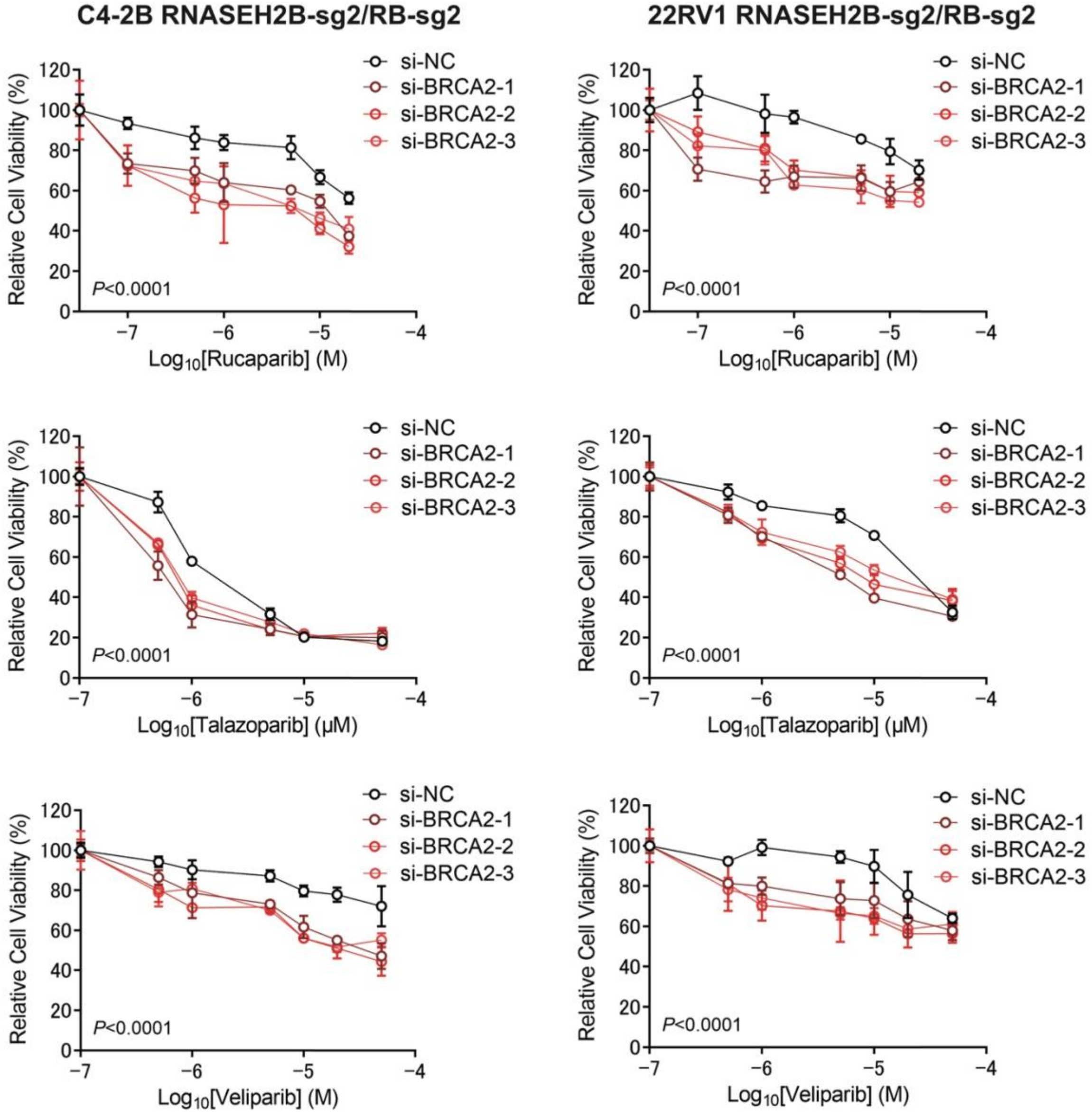


**Fig. S7. BRCA2 deletion re-sensitizes RNASEH2B/RB1 double gene knockout (DKO) cells to PARP inhibitors (PARPis).** BRCA2 was deleted in DKO C4-2B and 22RV1 cells using RNA interference with three different siRNAs. After 48 hours, BRCA2 knockdown cells and control cells were treated with talazoparib, rucaparib and veliparib as indicated for additional 7 days. Cell viability was measured using alamarBlue assay. *P*-value was determined by two-way ANOVA.

#### Table S1. Genes identified from CRISPR/Cas9 screens

| 79 common genes | Number of CRISPR screens |
|---|---|
| RNASEH2A | 5 |
| ATM | 5 |
| RNASEH2B | 5 |
| MUS81 | 5 |
| LIG1 | 5 |
| FANCA | 4 |
| RNASEH2C | 4 |
| CHD1L | 4 |
| BRCA1 | 4 |
| BARD1 | 4 |
| AUNIP | 4 |
| RAD51B | 4 |
| XRCC3 | 4 |
| EME1 | 3 |
| PPP1R8 | 3 |
| C19orf40 | 3 |
| PALB2 | 3 |
| RAD51C | 3 |
| FANCE | 3 |
| PSMC3IP | 3 |
| RAD51D | 3 |
| BRCA2 | 3 |
| TRAIP | 3 |
| RAD51 | 3 |
| SWI5 | 3 |
| PPP2R4 | 2 |
| CDK5 | 2 |
| TRAPPC4 | 2 |
| XKR7 | 2 |
| WDR48 | 2 |
| KDM8 | 2 |
| FANCM | 2 |
| ZNF574 | 2 |
| PHF12 | 2 |
| SACM1L | 2 |
| SF3B5 | 2 |
| DDX46 | 2 |
| SARS | 2 |
| NUP62 | 2 |
| SF3B3 | 2 |
| LRWD1 | 2 |
| ARGLU1 | 2 |
| CREM | 2 |
| SMC6 | 2 |

| | |
|---|---|
| C11orf30 | 2 |
| FANCD2 | 2 |
| HUS1 | 2 |
| PNKP | 2 |
| ZNF512B | 2 |
| COMMD1 | 2 |
| TSC1 | 2 |
| ANAPC2 | 2 |
| CHRAC1 | 2 |
| CENPW | 2 |
| TONSL | 2 |
| SRSF11 | 2 |
| RBBP8 | 2 |
| ATR | 2 |
| PGD | 2 |
| GTF2B | 2 |
| FANCC | 2 |
| CTDP1 | 2 |
| UBE2T | 2 |
| HJURP | 2 |
| XRCC2 | 2 |
| POLR2B | 2 |
| MRE11A | 2 |
| ESCO2 | 2 |
| TIPRL | 2 |
| URB1 | 2 |
| SNRNP200 | 2 |
| RNF168 | 2 |
| SPACA4 | 2 |
| HELLS | 2 |
| EPN1 | 2 |
| BRD8 | 2 |
| NBN | 2 |
| CHTF8 | 2 |
| XRCC1 | 2 |

#### Table S2. IC50 of olaparib

| Cell lines | LNCaP | C4-2B | 22RV1 | PC3 | DU145 |
|---|---|---|---|---|---|
| **AAVS1-sg1 (μM)** | 4.222 | 2.475 | 6.378 | 2.265 | 49.94 |
| **AAVS1-sg2 (μM)** | 4.585 | 2.926 | 4.968 | 2.255 | 55.36 |
| **RNASEH2B-sg1 (μM)** | 0.03039 | 0.02061 | 0.0383 | 0.5466 | 5.038 |
| **RNASEH2B-sg2 (μM)** | 0.004479 | 0.1612 | 0.07162 | 0.7888 | 8.001 |
| **AAVS1 Average** | 4.4035 | 2.7005 | 5.673 | 2.26 | 52.65 |
| **RNASEH2B-KO Average** | 0.0174345 | 0.090905 | 0.05496 | 0.6677 | 6.5195 |
| **Fold change** | 252.6 | 29.7 | 103.2 | 3.4 | 8.1 |

### Table S3. List of reagents

| Small molecule inhibitors | | |
|---|---|---|
| **Name** | **Company** | **Cat #** |
| VE-822 | Selleck Chemicals | S7102 |
| olaparib | Selleck Chemicals | S1060 |
| veliparib | Selleck Chemicals | S1004 |
| rucaparib | MedChemExpress | HY-10617 |
| talazoparib | MedChemExpress | HY-16106 |

| Antibodies | | |
|---|---|---|
| **Name** | **Company** | **Cat #** |
| PARP | Santa Cruz Technology | sc-7150 |
| β-Tubulin | Santa Cruz Technology | sc-80011 |
| β-Actin | Sigma-Aldrich | A5441 |
| normal rabbit IgG | Santa Cruz Technology | sc-2027 |
| E2F1 | Cell Signaling Technology | 3742 |
| RNASEH2B | Sigma-Aldrich | HPA040084 |
| RB1 | Cell Signaling Technology | 9309 |
| BRCA1 | Santa Cruz Technology | sc-6954 |
| BRCA2 | Cell Signaling Technology | 10741 |
| RAD51 | Abcam | ab133534 |
| phospho-Histone H2A.X | Millipore | 05-636 |
| phospho-CHK1 (Ser345) | Cell Signaling Technology | 2348 |
| CHK1 | Cell Signaling Technology | 2360 |

| siRNA sequences | | |
|---|---|---|
| Name | Company | Sequence (5' - 3') |
| siNC | Sigma-Aldrich | MISSION siRNA Universal Negative Control #2 (#SIC001) |
| siBRCA2 #1 | Sigma-Aldrich | SASI_Hs01_00121791 |
| siBRCA2 #2 | Sigma-Aldrich | SASI_Hs01_00121794 |
| siBRCA2 #3 | Sigma-Aldrich | CCGAUUACCUGUGUACCCU |
| siE2F1 #1 | Sigma-Aldrich | SASI_Hs01_00162220 |
| siE2F1 #2 | Sigma-Aldrich | SASI_Hs01_00162222 |

| RT-qPCR primer sequences | |
|---|---|
| Name | Sequence (5' - 3') |
| BRCA1-Forward | GACTGTTTATAGCTGTTGGAAG |
| BRCA1-Reverse | TTTTGGAAGTGTTTGCTACC |
| BRCA2-Forward | AATGTCAGACAAGCTCAAAG |
| BRCA2-Reverse | TCATGTATTTTTCAGGTGGC |
| RAD51-Forward | CAGATTGTATCTGAGGAAAGG |
| RAD51-Reverse | ATGATTCAGTCTTTGGCATC |
| GAPDH-Forward | GTCATGGGTGTGAACCATGAGA |
| GAPDH-Reverse | GGTCATGAGTCCTTCCACGATAC |

| guide RNA sequences | |
|---|---|
| Name | Sequence (5' - 3') |
| RNASEH2B-sg1: Forward | CACCGTCATAGGTTAATCAAACTG |
| RNASEH2B-sg1: Reverse | AAACCAGTTTGATTAACCTATGAC |
| RNASEH2B-sg2: Forward | CACCGAGTGGAGAAGCAGAAATAG |
| RNASEH2B-sg2: Reverse | AAACCTATTTCTGCTTCTCCACTC |
| RB1-sg1: Forward | CACCGTGCTCGCTCACCTGACGAG |
| RB1-sg1: Reverse | AAACCTCGTCAGGTGAGCGAGCAC |
| RB1-sg2: Forward | CACCGCACCTCGAACACCCAGGCG |
| RB1-sg2: Reverse | AAACCGCCTGGGTGTTCGAGGTGC |
| AAVS1-sg1: Forward | CACCGTCACCAATCCTGT |
| AAVS1-sg1: Reverse | AAACACAGGATTGGTGAC |
| AAVS1-sg2: Forward | CACCGGACTTCCCAGTGT |
| AAVS1-sg2: Reverse | AAACACACTGGGAAGTCC |
| BRCA2-sg1: Forward | CACCGAAAGCGATGATAAGGGCAG |
| BRCA2-sg1: Reverse | AAACCTGCCCTTATCATCGCTTTC |

| ChIP-qPCR primer sequences | |
|---|---|
| Name | Sequence (5'-3') |
| BRCA1 promoter-Forward | CTGTAATTCCCGCGCTTT |
| BRCA1 promoter-Reverse | CCTCCCATCCTCTGATTGTA |
| BRCA2 promoter-Forward | CCGCTTTATTCGGTCAGATAC |
| BRCA2 promoter-Reverse | GCGGGTATTTCTCAGTGTG |
| RAD51 promoter-Forward | CCAGAGACCGAGCCCTAA |
| RAD51 promoter-Reverse | GCTTACGCTCCACTTCTCTAC |
| Control region-Forward | AATGCTGGGCTTCCAAGGA |
| Control region-Reverse | GACCTTGGTGACTGTTGAGGAAAC |
